# A cautionary tale on the cost-effectiveness of collaborative AI in real-world medical applications

**DOI:** 10.1101/2024.05.27.596048

**Authors:** Lucia Innocenti, Sebastien Ourselin, Vicky Goh, Michela Antonelli, Marco Lorenzi

## Abstract

Federated learning (FL) has gained wide popularity as a collaborative learning paradigm enabling trustworthy AI in sensitive healthcare applications. Nevertheless, the practical implementation of FL presents technical and organizational challenges, as it generally requires complex communication infrastructures. In this context, consensus-based learning (CBL) may represent a promising collaborative learning alternative, thanks to the ability of combining local knowledge into a federated decision system, while potentially reducing deployment over-head. In this work we propose an extensive benchmark of the accuracy and cost-effectiveness of a panel of FL and CBL methods in a wide range of collaborative medical data analysis scenarios. Our results reveal that CBL is a cost-effective alternative to FL, providing comparable accuracy and significantly reducing training and communication costs. This study opens a novel perspective on the deployment of collaborative AI in real-world applications, whereas the adoption of cost-effective methods is instrumental to achieve sustainability and democratisation of AI by alleviating the need for extensive computational resources.

## 1 Introduction

The application of artificial intelligence (AI) in the field of biomedicine is witnessing a rapid expansion. Nonetheless, a significant gap still exists between the developments of AI for healthcare and their practical deployment in real-world scenarios [1]. While AI systems hold promises in revolutionizing healthcare, they often fail to achieve model generalizability when tested on data that differs from the one used for model training [2], limiting their applicability as clinical tools.

Bias is among the main factors affecting model generalizability since the variability observed in hospital data is generally affected by several factors, including local acquisition protocols and population demographics [3]. Differences between clinical populations with varying socioeconomic status or demographic characteristics such as age, and ethnicity can significantly impact AI model performance [4], and potentially lead to reduced performance when applied to data from underrepresented groups [5, 6]. To mitigate the impact of data bias it is therefore crucial to train models on datasets providing a complete representation of the natural variability that can be observed across hospitals, ideally gathered from multiple centers. The classical paradigm for multi-centric AI development consists of gathering data from different sites into shared repositories, where model training can be operated [7]. This paradigm, known as Centralized Learning, is an established setting for the development of AI in healthcare, as it allows training AI systems on large collections of data, addressing the problem of data-driven bias [8–11]. However, sharing patient information across health institutions raises important concerns over data governance and privacy [12]. Current regulations such as the European GDPR [13, 14] or the American CCPA [15] prescribe important limitations to data sharing practices, with a subsequent impact on the feasibility of the centralized learning paradigm in real-world applications [12, 16]. The need for using multiple data sources for training AI models while complying with data protection regulations encourages the adoption of collaborative learning (CL) strategies, in which the exchange of sensitive information is minimized. CL can be defined as a machine learning paradigm in which different entities can collaborate and jointly perform an analytical task without the need for sharing the original data [12, 17, 18]. To this end, CL relies on the sharing of data abstractions, which can, for instance, take the form of model parameters, global statistics, or predictions. CL is gaining popularity in AI applications due to its appealing promises for accuracy/privacy trade-off [1, 19].

Federated Learning (FL) [20] has been identified as a key CL paradigm [21], focusing on collaboratively optimizing model parameters across clients, each holding local datasets (Figure 1, left). In FL, each client shares model parameters partially trained on the respective local data. A central server aggregates the locally trained parameters to define a global model, which is subsequently re-transmitted to the clients and used to initialize a new round of partial training. This process is iterated for several rounds of local training and aggregation, until convergence. The successful integration of FL in healthcare applications is expected to deliver robust AI-based models that could handle and exploit data heterogeneity across hospitals while guaranteeing data governance [21, 22].

**Fig. 1:**
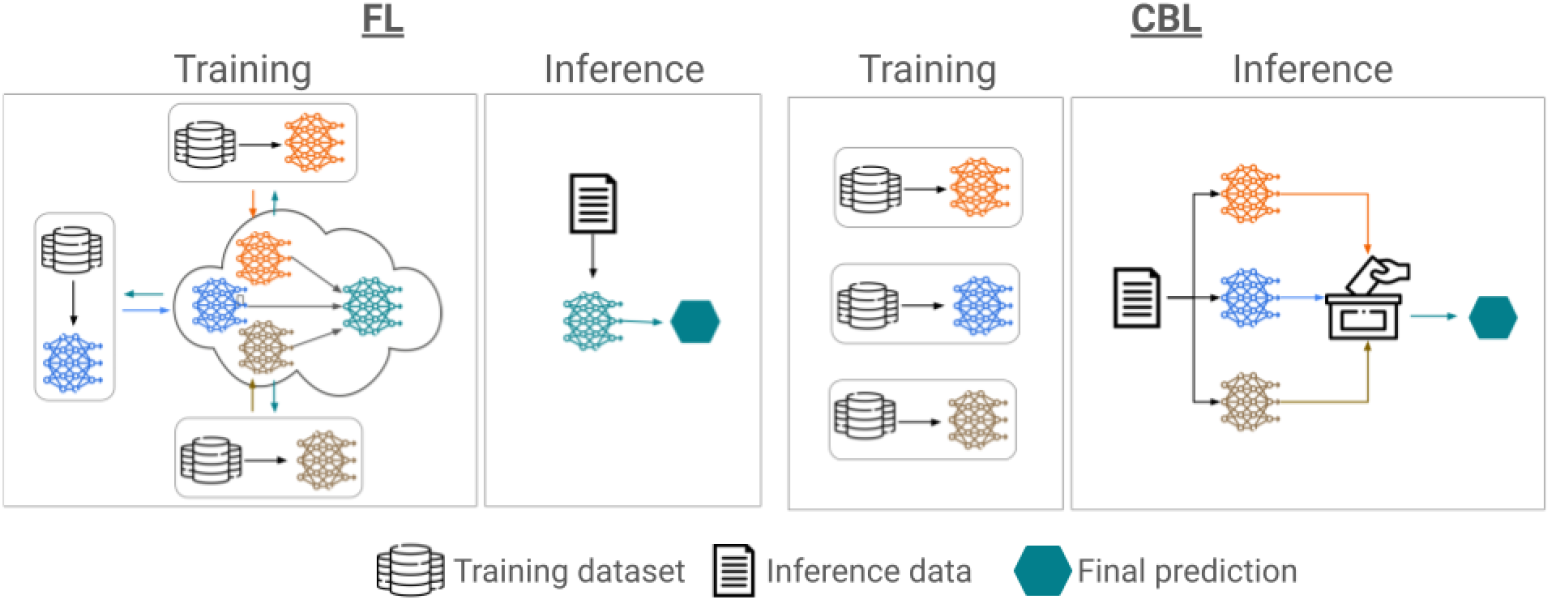
Training and inference phases for federated learning (FL, on the left) and consensus-based learning (CBL, on the right). In FL training is performed collaboratively to produce a common global model across clients. The global model is subsequently used for inference on new data instances. CBL instead requires clients to train a model on the respective local data independently. Inference on new data instances is performed collaboratively through consensus.

Although FL has significant advantages in security and efficiency, its implementation in real-world applications is not straightforward and can face technical and organizational challenges, reflecting the technology’s complexity. For example, performing distributed optimization across different clients requires coordinating the training process among different entities with varying availability, computational resources, and readiness level in performing AI tasks, which can be challenging in hospital settings [23]. Moreover, since typical distributed optimization approaches require multiple rounds of training and parameters exchange across parties, FL is usually associated with large communication costs and energy consumption [24].

Consensus-based learning (CBL) [25–27] may represent a valid CL alternative to FL. CBL does not rely on a shared training routine nor on a common model architecture across parties. Given a testing point, CBL combines the predictions obtained from the different models independently trained by each client on the local data. Therefore CBL relies on an off-line routine, in which information is exchanged only at inference time (Figure 1, right). Although ensembling methods have been widely used in medical imaging applications such as segmentation and classification tasks [28–38], their use in multi-centric CL setting is less popular.

Comparing the working schemes of FL and CBL of Figure 1, the main difference between FL and CBL is that FL relies on collaborative training, while CL on collaborative inference. While both paradigms aim for the same goal, the suitability of one setting over the other in real-world CL applications may vary substantially, as their deployment comes with different challenges. For example, FL typically requires several optimization rounds across clients and the setup of a dedicated communication infrastructure across hospitals. On the other hand, CBL requires the sharing of fully trained models, thus entailing privacy and intellectual property protection challenges. Bearing in mind the duality between these CL paradigms, when it comes to implementing collaborative frameworks in the real-world we currently lack quantitative benchmarks illustrating the quality and cost-effectiveness of specific CL settings in practical applications. To help promoting the implementation of collaborative AI in future multi-centric studies, in this study we introduce the first comparative analysis of the capabilities and cost-effectiveness of CBL and FL. The primary contributions of this work are:

i. To propose the first extensive head-to-head benchmark of FL and CBL for a variety of clinical tasks and data modalities.
ii. To compare a panel of FL and CBL methods from the state-of-the-art, with a particular focus on approaches suited to cope with data heterogeneity. To this end, we explore variants of CBL optimally integrating prediction uncertainty in the decision process.
iii. To identify heuristics for choosing the most appropriate CL approach based on the overall ability of each client to perform the specific task.

Our results show that in most of the evaluated benchmarks, CBL is a cost-effective alternative to FL, achieving comparable accuracy with the advantage of reducing training and communication costs. Our results open a novel perspective on the deployment of collaborative AI in the real-world, in which the opportune choice of CL paradigm and tasks can mitigate the implementation burden of CL.

In Section 2 we present the state of the art in CL and available benchmarks. Section 3 introduces the study design and the experiments. We discuss our results in Section 4, while detailing the adopted methodology in Section 5.

## 2 Collaborative learning benchmarks

Benchmarking is an essential step to assess the reliability of ML methods on specific tasks and datasets [39]. In healthcare, for example, initiatives such as MedPerf [40] are focusing on developing comprehensive platforms allowing researchers to test models on distributed data collections, focusing on medical imaging applications.

In the field of collaborative learning the commonly studied paradigm is FL, which has been mainly evaluated on publicly available datasets by using as metrics the model accuracy, convergence speed, and protection against cyber-attacks [12, 41–44], as well as communication overload and training time [45–47].

A common limitation in available FL benchmarks for healthcare is the partition of data among clients by artificial heuristics, a practice that often leads to an incorrect assessment of the actual performance of the methods studied. Works such as the FeTS challenge [16] and Flamby [48] highlight the importance of using natural partitions when we benchmark FL. In the FeTS challenge, researchers test different aggregation strategies for FL on a medical brain tumor segmentation dataset, collected from all around the world, while FLamby is a platform designed to compare various FL methods with centralized and locally trained models and allows researchers to add datasets or models to the benchmark. In Flamby, a variety of tasks and data typologies are considered, by including in the benchmark datasets from the literature. These datasets are split among clients using criteria such as the collection site and the acquisition method. Because of both their commitment to best representing the true characteristics of data and the focus on medical applications, FLamby and FeTS stand out from the variety of benchmarks for FL in the literature [41, 42, 49]. However, they also have two limitations: they do not compare FL with other collaborative paradigms and they don’t assess the cost-effectiveness of the various methods considered.

While few works benchmark FL in real-world scenarios for healthcare, there is even less literature on CBL for the same topic. To the best of our knowledge, the only studies testing CBL as a CL paradigm are Guha et al. (2019) [25], who applied CBL to publicly available datasets and compared it with centralized and local training, and Chaudhari et al. (2023) [50], who performed a comparison with FL on the ability to protect data from cyber-attacks. In the context of medical imaging applications, the capabilities of different CL paradigms were discussed in Gupta et al. (2023), without, however, providing any experimental evaluation [51].

Overall, current studies investigating CBL in collaborative healthcare applications are limited in scope and generalizability. A general reference point that allows comparison between CBL and FL paradigms while considering the accuracy and cost-effectiveness of these paradigms is still missing.

## 3 Results

The following section describes the benchmark’s design, introduces the CL methods and datasets used in the experiment, and presents the experimental results.

### 3.1 Study design

In this work, we used 7 different datasets, encompassing 3 tasks and 8 different data modalities, to benchmark 11 CL methods.

Table 1 illustrates the datasets used for the benchmark. A brief description of each dataset is available in subsection 5.1 (see Appendix A for more details).

**Table 1:** Datasets used for the benchmark. For each dataset, it is reported the number, the size of clients’ local datasets, the data modality, and the task along with the associated metric.

| Dataset | #Clients | Dataset size per client | Data Modality | Task [Metric] |
| --- | --- | --- | --- | --- |
| FedProstate | 6 | [32, 23, 27, 184, 5, 36] | T2 MRI | Segmentation [DiceScore] |
| FedHeart | 4 | [303, 261, 46, 130] | Tabular Data | Classification [Accuracy] |
| FedIXI | 3 | [311, 181, 74] | T1 MRI | Segmentation [DiceScore] |
| FedISIC | 6 | [12k, 3.9k, 3.3k, 225, 819, 439] | Dermoscopy images | Classification [Accuracy] |
| FedTCGA-BRCA | 6 | [311, 196, 206, 162, 162, 51] | Tabular Data | Survival [C-Index] |
| FedKiTS | 6 | [12, 14, 12, 12, 16, 30] | CT Scans | Segmentation [DiceScore] |
| FeTS | 23 | 4 to 511 (*Details in Appendix A*) | T1, T1CE, T2 & FLAIR MRIs | Segmentation [DiceScore] |

For each dataset, we compared the performances of the following paradigms: FL, CBL, centralized, and local learning. Local learning refers to the simple training independently performed by each client using only their local data, and centralized learning refers to training a model through the union of all the local datasets. The model obtained through centralized learning (*centralized model*) represents the upper limit of model accuracy expected for each dataset, while the model obtained through local learning (*local model*) provides the baseline accuracy that a single client could achieve on its own. Because the model trained with centralized learning sees all the data during the training, we expect it to generalize better than local models.

Concerning FL methods, our benchmark covers standard aggregation strategies already used in FL benchmarks for healthcare in the literature [41, 48]. We included the standard aggregation mechanism proposed in the seminal work of McMahan et al., FedAvg [20], along with subsequent approaches aimed at mitigating the impact of client heterogeneity in the optimization: FedProx,Scaffold, as well as FedAdam, FedYogi, and FedAdagrad [52–54]. Table 2 presents an overview of the 6 FL and 5 CBL methods used in the benchmark.

**Table 2:** Collaborative learning (CL) methods evaluated in the benchmark: six methods for federated learning (FL) and five methods for consensus-based learning (CBL).

| Method | CL Paradigm | Reference |
| --- | --- | --- |
| FEDAVG | FL | McMahan et. Al., 2017 [55] |
| FEDPROX | FL | Li et Al., 2018 [52] |
| SCAFFOLD | FL | Karimireddy et Al., 2020 [53] |
| FEDADAM | FL | Reddi et Al., 2020 [54] |
| FEDADAGRAD | FL | Reddi et Al., 2020 [54] |
| FEDYOGI | FL | Reddi et Al., 2020 [54] |
| AVG | CBL | Guha et Al., 2019 [25] |
| MV | CBL | Safdar et Al., 2021 [56] |
| STAPLE | CBL | Warfield et Al., 2004 [57] |
| UBE | CBL | *Inspired by* Ruta et Gabrys, 2000 [58] |
| ABE | CBL | *Inspired by* Ruta et Gabrys, 2000 [58] |

For CBL methods, we tested several classical fusion strategies to combine the predictions of local models, such as plain averaging (Avg) and majority voting (Mv). Moreover, for segmentation tasks, we used Staple, which optimizes consensus through expectation-maximization. Finally, we proposed two CBL methods based on decision averaging: uncertainty-based (Ube) and autoencoder-based ensembling (Abe). Both methods rely on the weighted average of local predictions. For Ube the weights are estimated by quantifying the uncertainty of the predicted labels. For Abe the uncertainty was quantified by the reconstruction error on the testing point of an autoencoder trained on the local dataset. This measure is a proxy for out-of-distribution detection [59].

Further details on all the methods are available in Section 5.2. Because of the characteristics of each fusion strategy, not all CBL methods can be applied to every task. Staple only applies to segmentation tasks, and the Mv aggregation is not compatible with the regression task of FedTGCA-BRCA.

We analyzed each collaborative method for its accuracy and cost-effectiveness. As a measure of task accuracy, the dice score, the balanced accuracy, and the C-Index were used, respectively, for segmentation, classification, and survival tasks, while we measured the cost-effectiveness by the *training time* and *bandwidth usage*. The training time was estimated as the total time necessary to obtain the collaborative model and the bandwidth as the total data transferred on the network during the training. We considered a scenario in which all clients participate in every round of FL training (no client selection), and we used the same model architecture across clients for each experiment. The performance of different CL methods was evaluated on the same cross-validation partitions.

### 3.2 Benchmark results

Figure 2 shows the mean accuracy, over folds, of each method across datasets grouped by paradigm: centralized learning, local learning, FL, and CBL. Given FeTS’s large number of clients (23), only aggregated statistics of the local models’ accuracies are reported in the figure (see Appendix A for results of each local model). The figure illustrates that collaborative learning generally outperforms local learning in terms of accuracy, achieving for some datasets accuracies nearly equivalent to those of centralized models, as observed with the FedTGCA-BRCA and FedHeart datasets. However, Figure 2 also indicates that no single collaborative learning paradigm consistently outperforms the others across all datasets. For example, for FedIXI (Figure 2e), FL performs worse than local learning, whereas CBL achieves higher accuracy. Conversely, for FedKiTS (Figure 2f), FL improves the local model prediction, while CBL degrades it.

**Fig. 2:**
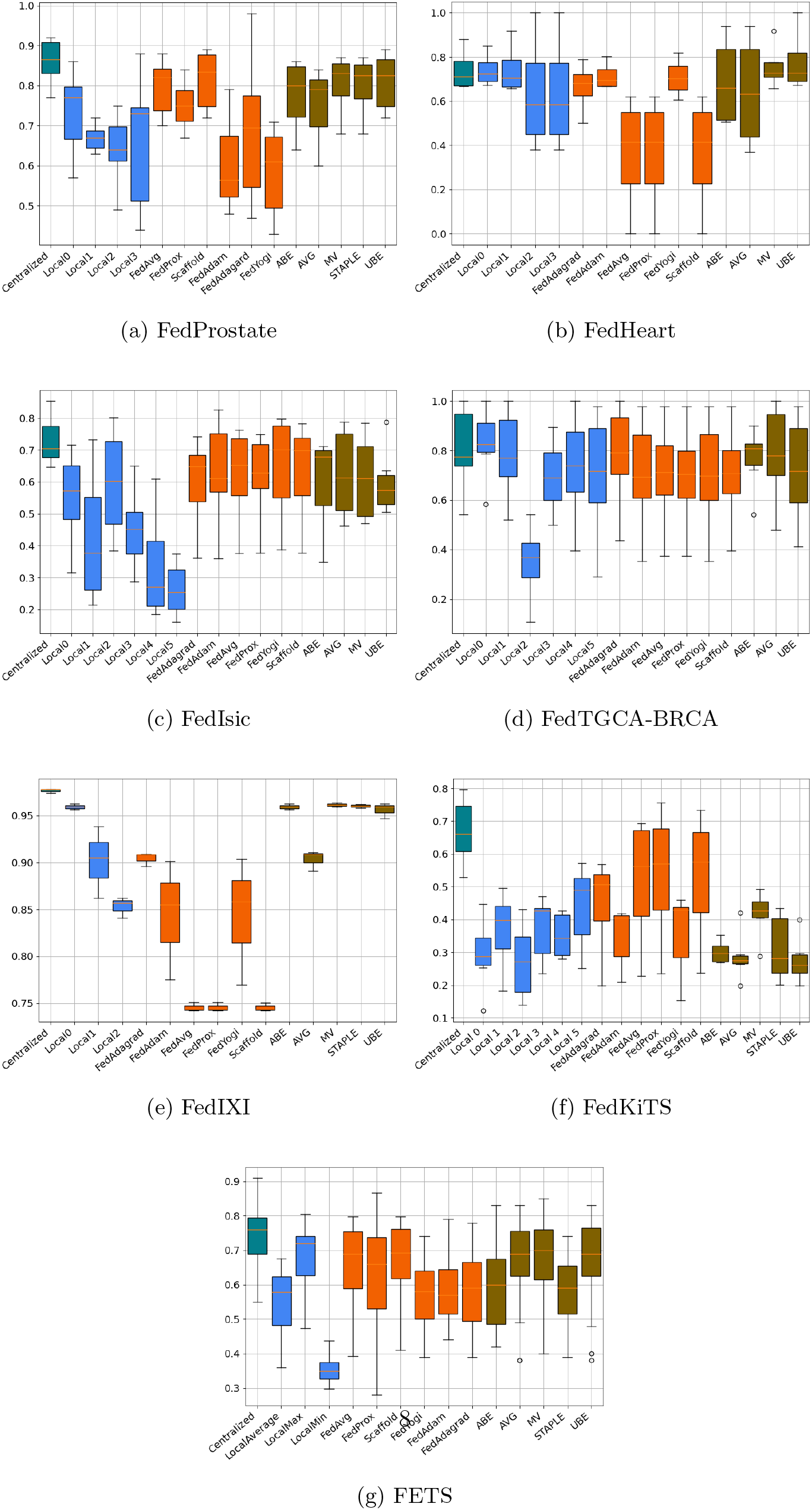
Results obtained by centralized learning (green), local learning (blue), federated learning (orange), and consensus-based learning (brown) methods. The boxplot represents the accuracy among test sets for centralized learning, local models, and CL methods. For FeTS, local accuracy results are aggregated due to the large number of clients.

Table 3 contains the summary results of the pair-wise Kruskal-Wallis test used to evaluate the difference in accuracy performance between FL and CBL methods along the various datasets. A significant difference was noted only in the FedIXI dataset (*p <* 0.05), with CBL performing better than FL. However, this significance did not hold after applying the Bonferroni correction for multiple comparisons across tasks.

**Table 3:** Summary of Kruskal-Wallis test used to assess the difference between accuracies of federated learning (FL) and consensus-based learning (CBL) methods along datasets. The only significant difference is observed in FedIXI (*p <* 0.05), where CBL is associated with better performances than FL. The statistical significance does not survive after Bonferroni correction for multiple comparisons across tasks.

| Experiment | F-statistic | P-Value |
| --- | --- | --- |
| FedProstate | 1.677 | 0.19 |
| FedHeart | 3.377 | 0.05 |
| FedIxi | 6.533 | **0.01\*** |
| FedIsic | 2.700 | 0.10 |
| FedTGCA-BRCA | 2.770 | 0.09 |
| FedKiTS | 3.266 | 0.07 |
| FeTS | 1.438 | 0.23 |

The Friedman test was applied to assess whether there was any statistically significant difference among the ranking of the methods in terms of testing accuracy among datasets, grouped according to the considered tasks (segmentation, regression, and classification). Significance was assessed by computing the *χ*^2^ statistic. We also compared the overall ranking of the methods across all tasks, by however excluding the method Staple (which applies only to segmentation tasks), and the task TGCA-BRCA (which does not support Mv as CBL method).

The results in Table 4 confirm that no statistical difference exists among the rankings of the methods, either when testing the tasks jointly or separately.

**Table 4:** Summary results for the Friedman test across tasks. No significant difference between the rankings of federated learning and consensus-based learning methods was identified across datasets for each specified task.

| Task | $\chi^2$ | p-value |
| --- | --- | --- |
| OVERALL | 8.039 | 0.624 |
| CLASSIFICATION | 6.07 | 0.62 |
| SEGMENTATION | 8.04 | 0.73 |
| SURVIVAL | 7.00 | 0.42 |

The numerical evaluation of the cost-effectiveness of CBL and FL, presented in Table 5, shows that the training of CBL methods is on average 8 to 30 times faster than the one required by the federated methods since CBL does not require multiple training rounds. For the same reason, the usage of bandwidth of CBL is from 35 to 120 times lower than for FL.

**Table 5:** Comparison of training time and bandwidth usage between federated learning (FL) and consensus-based learning (CBL). For both metrics, CBL is far less costly than FL.

|  | Training Time [min] |  |  | Bandwidth [MB] |  |  |
| --- | --- | --- | --- | --- | --- | --- |
|  | FL | CBL | FL Increment | FL | CBL | FL Increment |
| FedProstate | $1.2 \cdot 10^3$ | $1.1 \cdot 10^2$ | $\times 8$ | $4.0 \cdot 10^3$ | $1.4 \cdot 10^2$ | $\times 35$ |
| FedHeart | 0.1 | $2.0 \cdot 10^{-2}$ | $\times 9$ | $3.2 \cdot 10^1$ | $9.0 \cdot 10^{-2}$ | $\times 38$ |
| FedIxi | 5.5 | $6.4 \cdot 10^{-1}$ | $\times 8$ | $1.5 \cdot 10^2$ | 3.0 | $\times 50$ |
| FedIsic | $1.9 \cdot 10^2$ | $1.0 \cdot 10^1$ | $\times 18$ | $6.8 \cdot 10^3$ | $1.9 \cdot 10^2$ | $\times 58$ |
| FedTCGA | $4.5 \cdot 10^{-1}$ | $2 \cdot 10^{-2}$ | $\times 26$ | 0.9 | 5.42 | $\times 36$ |
| FedKiTS | $5.4 \cdot 10^1$ | $6.4 \cdot 1.5 \cdot 10^2$ | $\times 30$ | $8.8 \cdot 10^4$ | $7.4 \cdot 10^2$ | $\times 120$ |
| FeTS | $3.4 \cdot 10^4$ | $2.9 \cdot 10^3$ | $\times 12$ | $3.7 \cdot 10^4$ | $4.5 \cdot 10^2$ | $\times 83$ |

## 4 Discussion

Our results show that for typical medical data analysis tasks, CBL leads to performances at par with FL, albeit with a fraction of computation cost and bandwidth occupation. This result entails profound implications for the real-life adoption of AI in healthcare.

The key advantage of CBL relies on its asynchronous training approach, thus easing the collaboration among partners, as hospitals simply need to train their local models or apply existing ones to join the collaboration. Moreover, the withdrawal (or introduction) of a participant can be simply achieved by removing (or adding) their local model without the need for complex procedures [60]. The complexity of local models can also be adapted to the availability of local resources, thus mitigating the problem of hardware heterogeneity in federated setups [61, 62], and promoting the adoption of AI in healthcare. In FL, besides tuning the hyperparameters of the local models, it is also often necessary to tune method-specific hyperparameters, which increases the task complexity compared to CBL. Using the same hyperparameters on all FL methods can lead to variability in the accuracy across methods. In contrast, CBL methods exhibit lower variance among results.

While the generalization properties of CBL may improve by ensembling models trained on heterogeneous datasets [63, 64], it is known that the distributed optimization procedure of FL suffers from data heterogeneity, which causes degradation in convergence and accuracy [20, 53]. Such difference suggests that adopting CBL could be beneficial in collaborations among hospitals with heterogeneous data, reflecting for example varying geographical location or image acquisition techniques.

The CL paradigms presented in the benchmark allow the development of AI technologies compliant with restrictions on data sharing, albeit with some differences: the use of CBL requires the sharing of local models, thus possibly impacting the intellectual property (IP) of the model itself. On the other side, several studies are exploring techniques for FL where clients receive models tailored to their contribution [18], aiming at resolving related IP.

The analysis addressed in this paper focuses on the cost-effectiveness of CL to boost the deployment of AI in real-life healthcare applications. However, CL does not necessarily provide quantifiable privacy guarantees [65, 66]. Cryptographic tools such as secure aggregation (SA) [67, 68] or multi-party computation (MPC) [68], and differential privacy (DP) [69] are classical privacy-preserving methodologies proposed in the FL literature. While SA, MPC, and DP are widely investigated in FL, the analysis of CBL from a privacy-preserving perspective is less explored. The study [50] proposes the use of MPC for securely aggregating predictions in CBL, and in [27] a preliminary study compares the effectiveness of DP applied to FL and CBL. These studies show that privacy-preserving techniques can be adopted for CBL, with a potentially smaller impact on the final model accuracy when compared to FL.

With this work, we hope to raise awareness of the importance of cost-effective CL in the actual deployment of AI models on medical tasks. Starting from the empirical results here presented, future works should be devoted to developing quantitative measures of a CL system’s effectiveness before deploying a collaborative infrastructure. Such a theory would allow the identification of optimal collaborative paradigms based on accounting for several aspects, such as client availability, data heterogeneity, and security requirements. Moreover, in evaluating the cost-effectiveness, estimation of the CO2 emissions and economic cost of the training would bring added value to the analysis.

## 5 Methods

In this section, we detail the datasets and the CL paradigms adopted for our benchmark.

### 5.1 Datasets

The benchmark is composed of seven heterogeneous datasets representing real-world case studies. Five of the datasets (FedHeart, FedIXI, FedISIC, FedTGCA-BRCA, FedKiTS) have been benchmarked for FL in FLamby [48] and in our work we chose the network and loss coherently with their work. For FedProstate we used a 3D U-Net [70], as standard in the literature for segmentation problems, while for FeTS we trained a centralized model using both a 3D U-Net and a pre-trained SegResNet [71] as networks, and we used the latter as it achieved better performance. We evaluated the segmentation tasks by using the dice score (DSC). Further details on the composition of the datasets, the preprocessing steps, and the training task are provided in Appendix A.

#### FedProstate

We obtained this dataset by gathering data from 3 major publicly available datasets on prostate cancer imaging analysis (Medical Segmentation Decathlon [72], Promise12 [73], ProstateX [74]), and by a clinical dataset from the Guy St. Thomas Hospital, London, UK. The dataset contains T2-magnetic resonance imaging (MRI) and segmentation masks of the whole prostate. We used the acquisition protocol (MRI with or without endorectal coil) and the scanner manufacturer as splitting criteria. In this way, we obtained a dataset with 6 clients: 4 of them we used for the training, and 2 we kept as independent test sets.

#### FedHeart

Firstly published in a centralized version [75] this dataset contains tabular information about 740 patients from 4 hospitals. The dataset is composed of 13 features: age, sex, chest pain type, resting blood pressure, serum cholesterol, blood sugar, resting electrocardiographic results, maximum heart rate, exercise-induced angina, ST depression induced by exercise, slope of the peak ST segment, number of major vessels, and thalassemia background. In [48] a multi-centric version of the dataset was proposed by allocating to each of the 4 clients the data from one of the hospitals. In this benchmark, we trained a fully connected ReLU network to solve the task of binary classification (presence or not of heart disease), and we used the classification accuracy as the evaluation metric.

#### FedIXI

We considered the brain imaging dataset presented in [48]. The dataset comprises brain T1 MRIs from 566 patients from the IXI dataset [76]. In Flamby the data was partitioned into 3 groups, based on the 3 hospitals where MRI acquisitions were performed. The task here considered is brain segmentation based on the training of a 3D U-Net [70] taking as input the patient’s T1 MRI and returning the brain segmentation mask. The loss adopted to train the network was the DSC.

#### FedISIC

The data for the ISIC dataset were originally published by the International Skin Imaging Collaboration [77], which collects dermoscopy images of different types of skin cancer. The challenge’s task is skin disease classification among 8 possible classes. The multi-centric version of this dataset - obtained using as splitting criteria the imaging acquisition system - is made of 6 clients, and characterized by elevated data heterogeneity in size, feature, and label distribution. We trained an EfficientNet [78] to solve the classification task and evaluated it through classification accuracy.

#### FedTCGA-BRCA

The original TCGA-GDC dataset [79] is a large collection of data from cancer genomic studies. In [48], they proposed a multi-centric version of TCGA-GDC by considering tabular data with 39 input features representing the patient’s demographic and medical condition, and the patient survival probability as a task. The collaboration is composed of 6 clients, corresponding to the data acquisition site, and we trained a fully connected LeakyReLU network, using the G-Index as the evaluation metric.

#### FedKITS

The KiTS dataset contains CT scans and segmentation masks for both the kidney and the tumor. The original version was proposed for the KiTS19 Challenge [80], and a collaborative version with 6 clients was defined in [48] based on the hospital producing the data. As a model, we used a 3D U-Net from the nnU-Net library [81], and the metric is DSC.

#### FeTS

This dataset was published in the context of the FeTS Challenge [82]. It is a multi-modal dataset, using 4 different brain MRI modalities to provide a multi-class segmentation among the different regions of gliomas. The data is partitioned based on the 23 data acquisition sites. Clients have variable sizes: the smallest has only 4 data points, the largest 511. We used a pre-trained SegResNet [71], and we evaluated it by the average DSC among different classes.

### 5.2 Collaborative Learning Methods

Let’s consider a collaborative setting with *M* clients. A dataset 𝒟 belonging to client *i* is composed by data samples 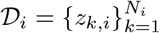, being *N*_*i*_ the dataset size. We consider a model *f* with parameters *θ*, a loss function ℒ, and we denote the prediction of a data instance *z* by *h* = *f* (*z, θ*).

**FL** is a collaborative paradigm associated with the optimization of a loss distributed among *M* clients:

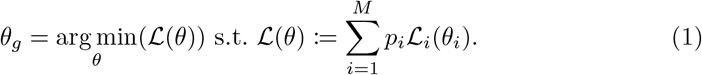

In Eq. (1), the losses of local models (*θ*_*i*_) are averaged by using the weights *p*_*i*_, and the obtained loss is minimized and leads to the global model *θ*_*g*_.

To solve the problem (1) different optimization strategies have been proposed in the past, aiming at mitigating critical distributed optimization challenges induced for example by the problem of data heterogeneity across clients. We consider here a comprehensive panel of state-of-the-art optimization approaches, which are at the core of the FL literature:

- **FedAvg**[55] is the backbone of FL optimization, and is based on an iterative process where, at each optimization round *r*, clients execute a fixed number of local stochastic gradient descent steps and send the partially optimized model 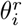 to the server. The server averages the received models according to the weights *p*_*i*_ to obtain a global one, 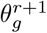. The global model is then sent to the clients to initialize the next optimization round.
- **FedProx**[52] tackles the problem of federated optimization with data heterogeneity across clients. This approach extends FedAvg by introducing a proximal term to the local objective function to penalize model drift from the global optimization during local training. The proximal term is controlled by a trade-off hyperparameter, *µ*. Being *r* the current round, the optimization problem becomes:

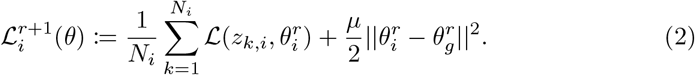
- **SCAFFOLD[53]** addresses the limitations of FedAvg in scenarios with heterogeneous (non-iid) data by utilizing control variates, which effectively reduces variance and corrects for client-drift in the local updates. To achieve this, SCAFFOLD maintains a state for each client (client control variate) and the server (server control variate).
- **FedAdam, FedYogi, FedAdagrad[54]** are adaptations of Adam[83], Yogi[84], and Adagrad[85] optimizers, designed to suit FL’s decentralized optimization setup.

While many additional optimization approaches can be found in the FL literature [24], they are often based on extensions of the methods here presented, for example focusing on different federated loss functions [86, 87], aggregation strategies [88–90], or communication strategies [91, 92].

**CBL** is a class of machine learning algorithms relying on the well-explored concept of ensembling [53, 93, 94]. The rationale of CBL consists of obtaining robust predictions by aggregating decisions obtained by independently trained weak predictors.

The CBL paradigm is based on a *training* phase in which *M* models *f* (·, *θ*_*i*_) are independently trained on separated data collections 𝒟_*i*_, by minimizing local objective functions ℒ_*i*_. These models are subsequently collected and, for a given test data *z*^*′*^ at *inference* time, the predictions *h*_*i*_(*z*) = *f* (*z, θ*_*i*_) from all the clients’ models are computed and aggregated by applying an ensembling (or fusion) strategy [95]:

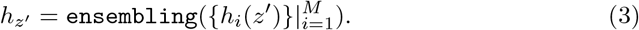

Typical ensembling methods proposed in the literature are:

**Majority voting** (Mv) [56], often used in classification tasks, aggregates predictions by selecting the most commonly predicted class among the experts.

**Staple** [57], which is an expectation-maximization algorithm that, iteratively, first computes a weighted average of each local prediction, and then assigns a performance level to each client’s segmentation, which will be used as weights for the next step.

**Decision averaging** (DA) [58, 96], an approach based on probabilistic principles: given different datasets 𝒟_1_, … 𝒟_*M*_ and relative local models *f* (·, *θ*_1_), … *f* (·, *θ*_*M*_), ensembling is obtained as as

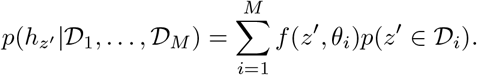

Different CBL algorithms can be obtained based on the estimation of the probability *p*(*z*^*′*^ ∈𝒟_*i*_), defining the relative weight of the local expert in the definition of the consensus. In this work, among the possible DA algorithms, we consider the following:

- **Average** (Avg) approximates *p*(*z*^*′*^∈𝒟_*i*_) as a uniform distribution. While this strategy can be adopted in all tasks, for classification and segmentation problems we consider the distribution probability obtained through a *softmax* function.
- **Uncertainty based ensembling** (Ube) in which the probability *p*(*z*^*′*^∈𝒟_*i*_) is approximated as the uncertainty of the local model on the prediction task. Ube defines averaging weights based on the model uncertainty quantified by the total element-wise variance at inference time:

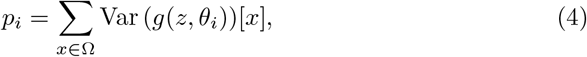

where Ω is the set of voxels in *z*, Var (·)[*x*] is the sampling variance estimated from *S* stochastic forward passes of the model, computed at voxel *x*. Through a softmax function, we define averaging weights:

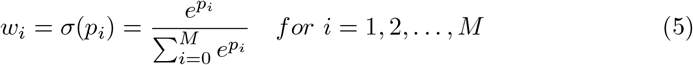
- **Autoencoder based ensembling** (Abe) computes a proxy for the probability *p*(*z*^*′*^ ∈ 𝒟_*i*_) by modeling the variability of the local dataset through autoencoders trained by each client on the respective local dataset. At inference time, weights are defined as the reconstruction error *e*_*i*_ on the testing data point *z*.

### 5.3 Experimental Setting

For each dataset in the benchmark, we split local partitions into training and testing. For FeTS and FedProstate, we used respectively a 4-fold and a 5-fold splitting approach, while for the other datasets, following [48], we ran all the experiments three times with different seeds. The obtained results are the average among the runs.

For FL, a federated infrastructure was simulated using FedBioMed version 4.1^1^ [97], an open-source framework for federated learning. For each FL method, the final global model was collected, along with the local models independently trained by each client. The local models are used to estimate the non-collaborative model performance and for generating the predictions subsequently aggregated with the CBL methods. As upper-bound for the comparison, a centralized model has been trained by pooling together all the local training sets.

To make the comparison as fair as possible, for each dataset we fixed the total number of training steps to be executed in each experiment. Specifically, a number *E* of epochs is defined a priori, corresponding to the number of training epochs executed by the centralized model. For the local training, each client executes 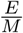 epochs, being *M* the number of clients in that configuration. For the federated strategies, we finetuned the number of local SGD steps *s* executed at each round, while the number of rounds is defined as follows: *R* = *E*·*N*_*T*_ */M/B/s*, where *B* is the batch size and *N*_*T*_ is the total number of samples in the training set.

All the experiments were developed in Python, version 3.7.6, using the following libraries: PyTorch version 2.0.1 [98] and Monai 1.2.0 [99] for the model architecture and torchio [100] version 0.19.1 with simpleITK [101–103] version 2.2.1 for the medical images management.

## Acknowledgements

LI and ML are supported by the 3IA Côte d’Azur Investments in the Future project managed by the National Research Agency (ref.n ANR-15-IDEX-01 and ANR-19-P3IA-0002). ML is funded by the grants (TRAIN - ANR-22-FAI1-0003) and (Fed-BioMed - ANR-19-CE45-0006).

## Appendix A Details on Federated Datasets, Architectures, and Hyperparameters

### A.1 FedProstate

The FedProstate dataset is the federated version of the 3 major publicly available datasets on prostate cancer imaging analysis, and of 1 private dataset:

- **Medical Segmentation Decathlon - Prostate** [72] provides 32 prostate MRIs for training.
- **Promise12** [73] consists of 50 training cases obtained with different scanners. Of those, 27 cases were acquired by using an endorectal coil.
- **ProstateX** [74] contains prostate MRIs acquired by using two different scanners (Skyra and Triotim, both from Siemens). Segmentations of 194 cases are available [104].
- **Guy St. Thomas Hospital dataset (King’s College London)**. This dataset is composed by 36 MRIs acquired during the clinical routine for patients with prostate cancer under active surveillance treatment. Images were acquired with a Siemens Aera scanner and an expert radiologist produced masks of the whole prostate gland. This dataset is used as an independent test set.

Datasets were split as in Table A1, to define centers characterized by specific image acquisition properties, thus allowing to obtain heterogeneous image distributions among centers. The common preprocessing pipeline applied to all the data comprised of flipping, cropping/padding to the same dimension, and intensity normalization. N4-bias-correction has also been applied to the data from Promise12 to compensate for the intensity artifacts introduced by the endorectal coil. Figure A1 shows an example of the resulting splits.

**Table A1:** FedProstate. Description of the different centers here considered for the distributed learning scenario, derived by partitioning the four datasets Decathlon, ProstateX, Promise12, and PrivateDS.

| ID | #Samples | Dataset | Subset Selection | Splitting Strategy |
| --- | --- | --- | --- | --- |
| Local0 | 32 | Decathlon | Full Dataset | 5-fold CV |
| Local1 | 23 | Promise12 | No Endorectal Coil | 5-fold CV |
| Local2 | 27 | Promise12 | Only Endorectal Coil | 5-fold CV |
| Local3 | 184 | ProstateX | Only Scanner Skyra | 5-fold CV |
| Local4 | 5 | ProstateX | Only Scanner Triotim | External test set |
| Local5 | 36 | PrivateDS | Full Dataset | External test set |

**Fig. A1:**
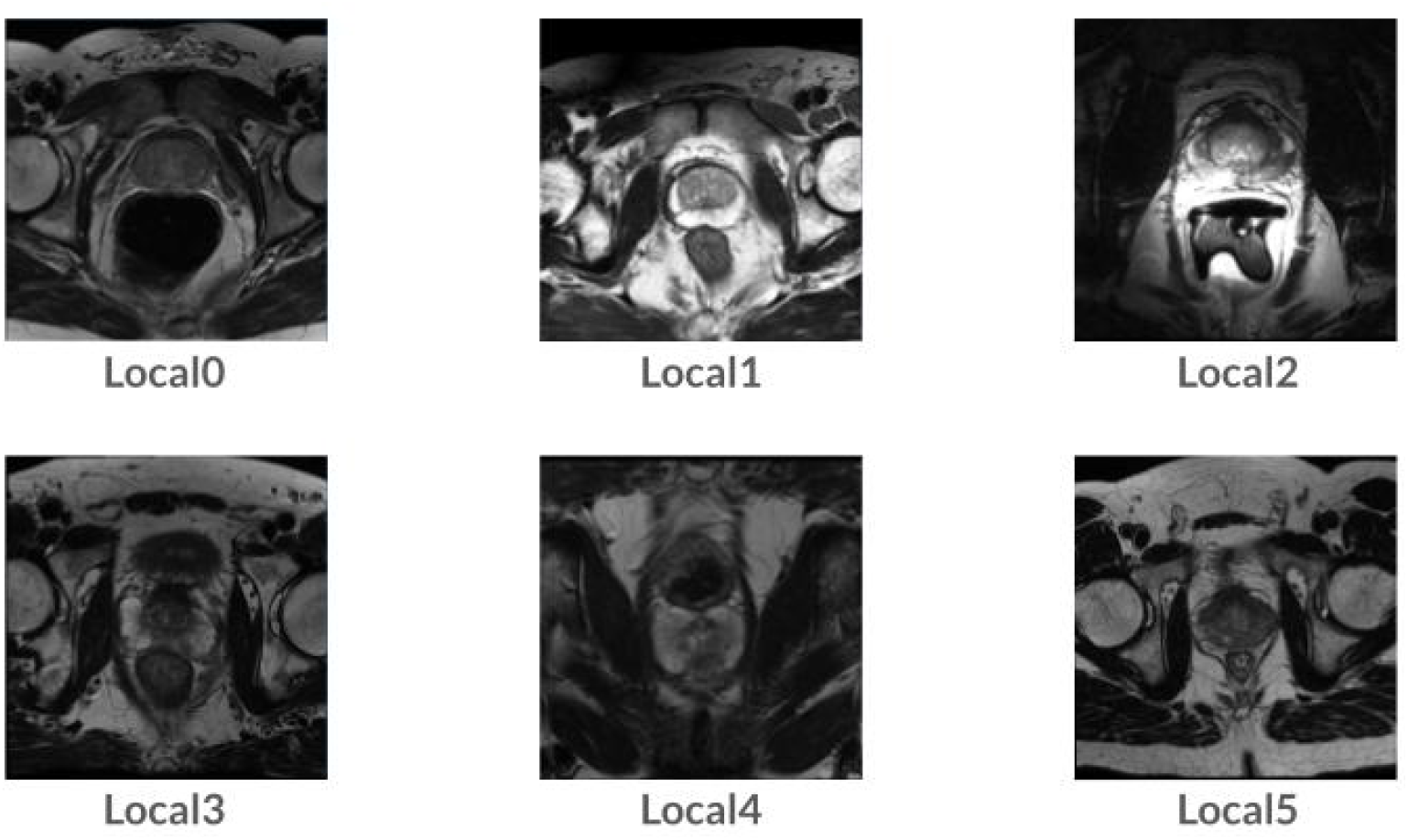
Examples of images from the FedProstate dataset showing the heterogeneity among different clients.

**Table A2:** FedProstate. Hyperparameters and respective values explored during the tuning phase. Selected value in **bold**. The selection of dropout value was driven by the need to use it for the UBE method. In red, the values selected for FedAdam, FedYogi and FedAdagrad, for which a different tuning was required.

| Parameter | Values |
| --- | --- |
| Learning Rate | 0.0001; **0.001**; 0.01; 0.1; 1 |
| Batch Size | 4, **8**, 16 |
| Dropout | 0.1, **0.3**, 0.5 |
| Local Steps | 10, 15, **20**, 25 |

#### Architectures and Parameters

The segmentation problem for this dataset was addressed by training a 3D UNet architecture with residual connections [70]. The training was based on optimizing the DSC, by using the AdamW optimizer for all experiments [105] but for SCAFFOLD, for which we used an SGD optimizer. The UNet implementation is available in the MONAI library^2^. The hyperparameters used for the training phase are available in the Table A2

### A.2 FedHeart

The dataset is a collection of tabular data, and the task consists of binary classification to recognize heart disease. In FLamby [48], a federated version has been proposed by using as subset selection criteria the hospital that provided the data. There are four hospitals (Cleveland’s, Hungarian, Switzerland, and Long Beach), so four clients. The preprocessing in FLamby has been done by removing missing values and encoding non-binary categorical variables as dummy variables.

#### Architectures and Parameters

A fully connected ReLU network with one hidden layer was used as a classification model. The training was based on optimizing the cross-entropy loss, with an AdamW optimizer. The details on the hyperparameters used for the centralized and local training are available in Table A3.

**Table A3:** FedHeart. Hyperparameters and respective values explored during the tuning phase. Selected value in **bold**. The selection of dropout value was driven by the need to use it for the UBE method.

| Parameter | Values |
| --- | --- |
| Learning Rate | 0.0001; 0.001; 0.01; **0.1**; 1 |
| Batch Size | **8**; 16; 32; 64 |
| Dropout | 0.1; **0.2**; 0.3 |
| Local Steps | 5; **10**; 20 |
| Centralized epochs | 10; **50**; 100 |

### A.3 FedIXI

The dataset contains T1 and T2 brain MRIs, as well as brain segmentation masks. In FLamby, the federation is obtained by using the hospital as a subset selection criteria, comprising three clients. Pre-processing pipeline comprehends volume resizing to 48 × 60 × 48 voxels, and sample-wise intensity normalization.

**Table A4:** FedIXI. Hyperparameters and respective values explored during the tuning phase. Selected value in **bold**. The selection of dropout value was driven by the need to use it for the UBE method.

| Parameter | Values |
| --- | --- |
| Learning Rate | 0.0001; **0.001**; 0.01; 0.1; 1 |
| Batch Size | **2**; 4; 6 |
| Dropout | 0.1; **0.2**; 0.3 |
| Local Steps | 5; **10**; 20 |
| Centralized epochs | **10**; 20; 30 |

#### Architectures and Parameters

The network used as a baseline model for this problem was a 3D UNet [70]. The training was based on optimizing the DICE Loss, by using the AdamW optimizer. The details on the hyperparameters used for the centralized and local training are available in Table A4.

### A.4 FedISIC

This dataset represents a skin cancer detection problem through image classification of CT scans. There are 8 different classes, with high distribution imbalance. Starting from the data available in the ISIC2019 dataset [77] the authors of FLamby obtained a federated dataset by allocating to a different client data obtained by using a different scan, so obtaining 6 different clients for the federation. The applied preprocessing is described in [106].

#### Architectures and Parameters

The network used as a baseline model for this problem was EfficientNet [78], as in FLamby. The training was based on optimizing the weighted focal loss, by using the AdamW optimizer. The details on the hyperparameters used for the centralized and local training are available in Table A5. When choosing the batch size, we applied different values for different local clients to account for the high variance in dataset dimensions. The final values are Local 0 and Local 1: 128, Local 2: 64, Local 3, Local 4 and Local 5: 32.

**Table A5:** FedISIC. Hyperparameters and respective values explored during the tuning phase. Selected value in **bold**. The selection of dropout value was driven by the need to use it for the UBE method.

| Parameter | Values |
| --- | --- |
| Learning Rate | **0.0005**; 0.005 0.001; |
| Batch Size | 32; 64; **128**; 256 |
| Dropout | 0.1; 0.2; **0.3** |
| Local Steps | 5; **10**; 20 |
| Centralized epochs | 50; 100; **150** |

### A.5 FedTCGA-BRCA

This dataset is composed of data from the TCGA-GDC portal, specifically those belonging to the breast cancer study, which includes features gathered from 1066 patients. The federated version is obtained by splitting the original dataset into 6 subsets, one for each extraction site, grouped into geographic regions. The task consists of predicting survival outcomes based on the patients’ tabular data, with the event to predict death. Each patient is defined by 38 features.

#### Architectures and Parameters

As a baseline, we used a fully connected LeakyReLU network. The training was based on optimizing the weighted focal loss, by using the AdamW optimizer. The details on the hyperparameters used for the centralized and local training are available in Table A6.

**Table A6:**
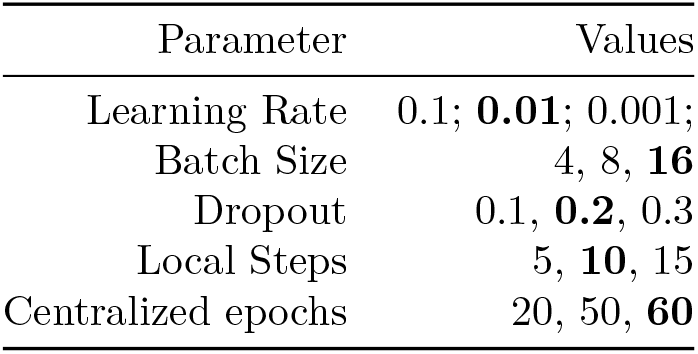
FedTCGA-BRCA. Hyperparameters and respective values explored during the tuning phase. Selected value in **bold**. The selection of dropout value was driven by the need to use it for the UBE method.

| Parameter | Values |
| --- | --- |
| Learning Rate | 0.1; **0.01**; 0.001; |
| Batch Size | 4, 8, **16** |
| Dropout | 0.1, **0.2**, 0.3 |
| Local Steps | 5, **10**, 15 |
| Centralized epochs | 20, 50, **60** |

### A.6 FedKiTS

The KiTS19 dataset is part of the Kidney Tumor Segmentation Challenge 2019 and contains CT scans of 210 patients along with the segmentation masks from 79 hospitals. In FLamby, the federated dataset is defined by splitting scans among different clients based on the providing hospital; they extracted a 6-client federated version by removing hospitals with less than 10 training samples. The preprocessing pipeline comprises intensity clipping followed by intensity normalization and resampling of all the cases to a common voxel spacing.

#### Architectures and Parameters

As a baseline, following FLamby implementation, a nnUNet [81] was used. The training was based on the optimization of the DICE Losse, by using the AdamW optimizer. The details on the hyperparameters used for the centralized and local training are available in Table A7.

### A.7 FeTS

The data are gathered from the FeTS 2022 Challenge. A more specific data description is available at the url ^3^. The dataset contains 1251 instances. Following the guidelines, we used a natural partitioning by the institution, obtaining a federation with 23 clients, for which the dataset size is available in Figure A2.

**Table A7:** FedKiTS. Hyperparameters and respective values explored during the tuning phase. Selected value in **bold**.

| Parameter | Values |
| --- | --- |
| Learning Rate | 0.1; **0.01**; 0.001; |
| Batch Size | 2, **4** |
| Dropout | 0.1, **0.2**, 0.3 |
| Local Steps | 30, 50, **100** |
| Centralized epochs | 2000, **5000**, 8000 |
| Optimizer | Adam, **AdamW**, SGD |

For each patient in the study, the following data modalities are available: native (T1), post-contrast T1-weighted (T1Gd), T2-weighted, and Fluid Attenuated Inversion Recovery (T2-FLAIR) volumes. An example of data for one patient is available in Figure A3.

Data in this FD are very heterogeneous, being acquired with different clinical protocols and various scanners from multiple data-contributing institutions.

The task is multi-class 3D image segmentation, and the labels are the GD-enhancing tumor (ET — label 4), the peritumoral edematous/invaded tissue (ED — label 2), and the necrotic tumor core (NCR — label 1). We segmented single sub-regions, not into intersections as was proposed in the FeTS challenge.

#### Architectures and Parameters

The network used is a SegResNet [71], which takes as input the multi-modal data and produces a segmentation mask. The model was trained by optimizing a DICE loss by an AdamW optimizer. As a preprocessing step, each data has been cropped to a common shape of 240×240×128 and intensity normalization has been applied. The details on the hyperparameters used for the centralized and local training are available in Table A8.

**Table A8:** FeTS. Hyperparameters and respective values explored during the tuning phase. Selected value in **bold**.

| Parameter | Values |
| --- | --- |
| Learning Rate | 0.1; 0.01; **0.001**; |
| Batch Size | 1, 2, **4** |
| Dropout | 0.1, 0.2, **0.3** |
| Local Steps | 1, **6**, 20 |
| Centralized epochs | 15, **30**, 50 |

**Fig. A2:**
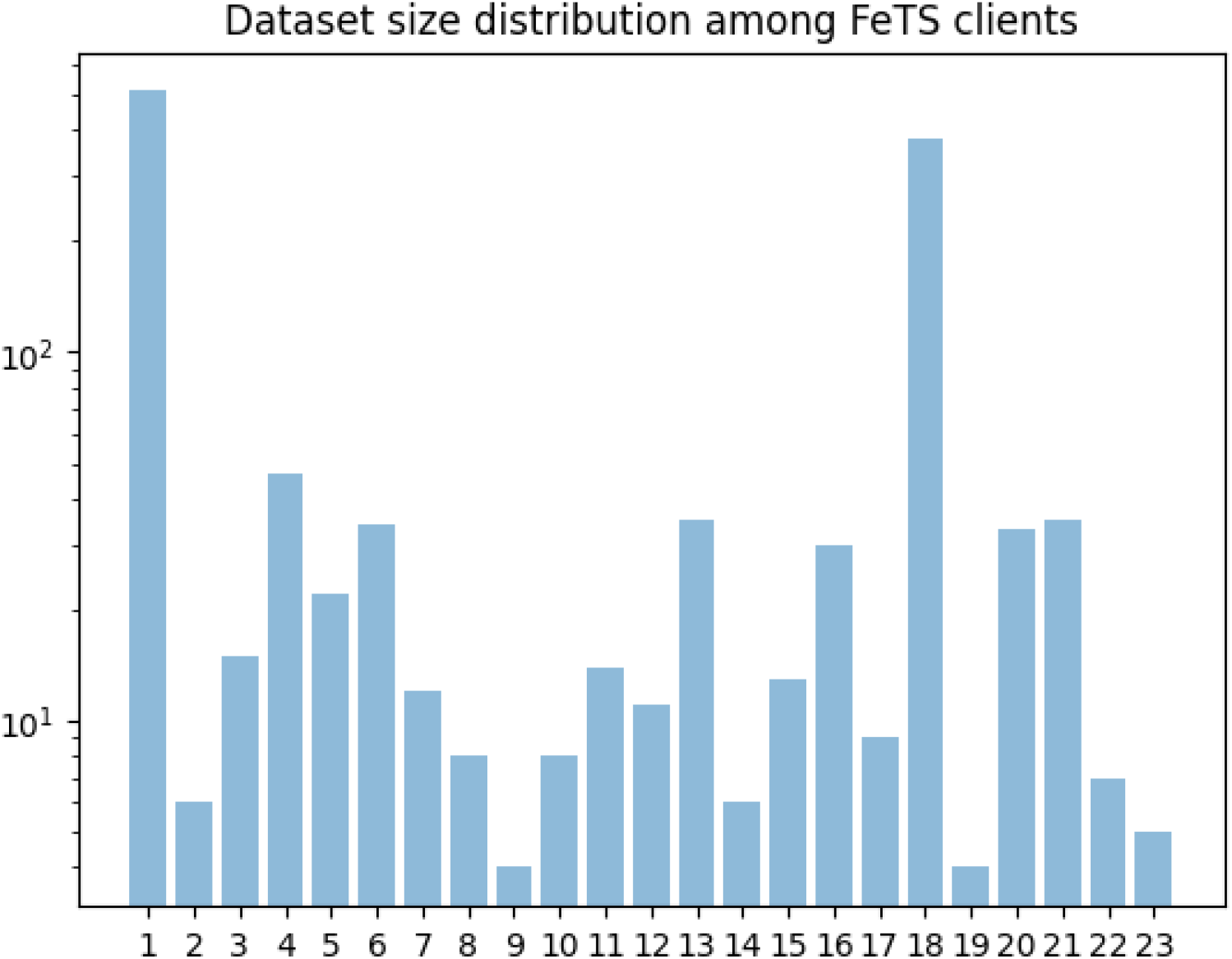
Distributions of dataset size among all the clients in the FeTS dataset

#### Results on local clients

Figure A4 shows the average DSC across splits and runs for the FeTS dataset. This figure completes Figure 2g.

## Appendix B Additional results

The Friedman test is a nonparametric method for evaluating the significance of differences between multiple classification algorithms against multiple datasets, comparing how well each model ranks (in terms of accuracy) across different datasets. In Section 3.2 we have shown the numerical results of the Friedman test, while in Figure B5 we show an analysis of the rankings on the various methods. We can read the figure as the representation of the probability of each method of ranking at a given position when tested on a dataset. The sparsity of the heatmap qualitatively suggests that no method emerges above the others to be systematically the best. These results are consistent with those of the statistical analysis of the p-values.

**Fig. A3:**
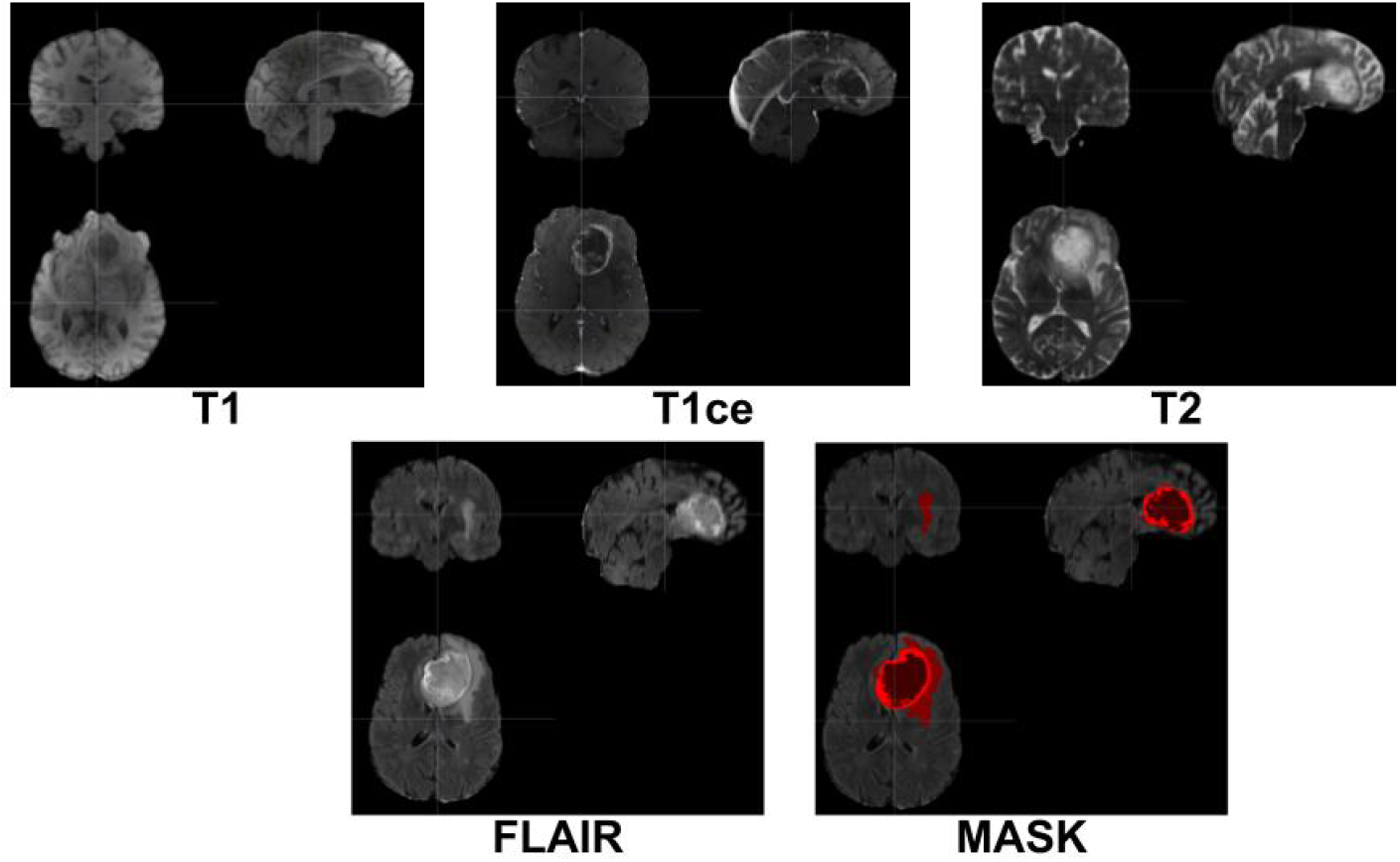
Examples of different modalities for one patient in the FeTS dataset. MASK represents the segmentation mask, which is used as ground truth for our segmentation problem.

**Fig. A4:**
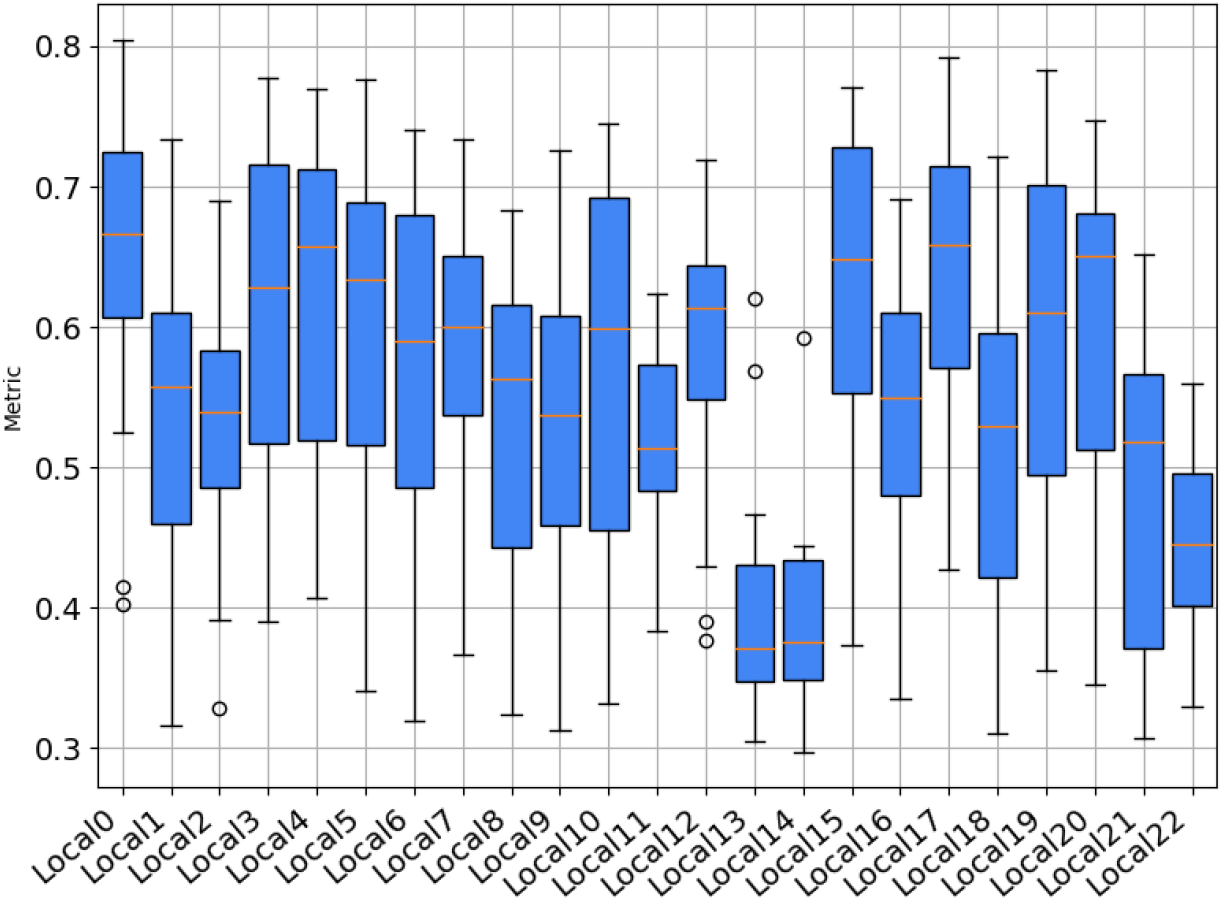
Average dice score obtained by locally-trained models on the FeTS dataset

**Fig. B5:**
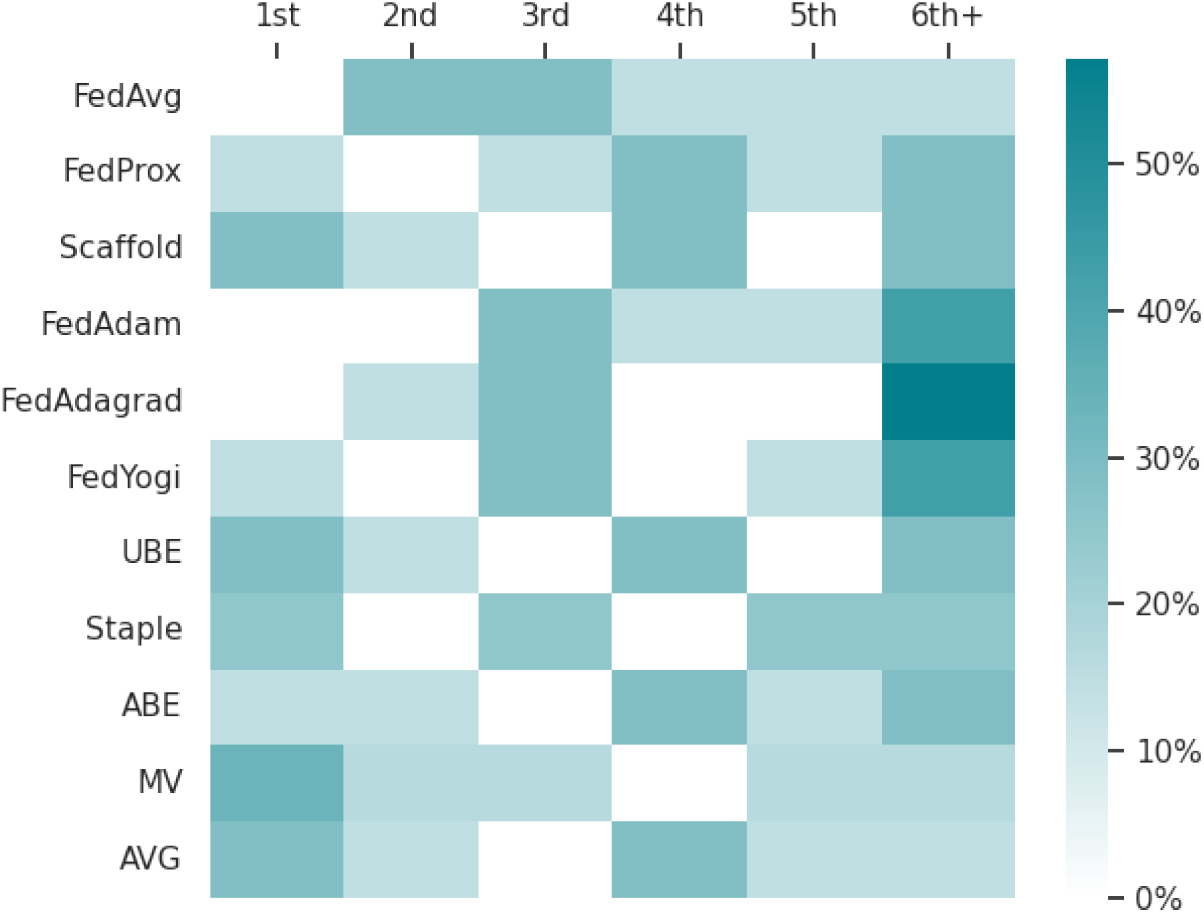
Performance ranking. Columns represent the ranking position for the accuracy of the methods across datasets. We note that not all the methods can be applied to each dataset: Staple can only be applied to segmentation tasks, and MV does not apply to the survival task of FedTGCA-BRCA. Overall, no method stands out in terms of overall best performance (Friedman test on segmentation (*p* = 0.73), classification (*p* = 0.62), and survival analysis (*p* = 0.42) tasks).

## Footnotes

1 https://fedbiomed.org/

2 https://monai.io/index.html

3 https://www.synapse.org/#!Synapse:syn28546456/wiki/617246

## References

[1] Collaborative learning without sharing data. Nature Machine Intelligence 3(459) (2021)

[2] Kelly, C.J., Karthikesalingam, A., Suleyman, M., Corrado, G., King, D.: Key challenges for delivering clinical impact with artificial intelligence. BMC medicine 17, 1–9 (2019)

[3] Singh, H., Mhasawade, V., Chunara, R.: Generalizability challenges of mortality risk prediction models: A retrospective analysis on a multi-center database. PLOS Digital Health 1(4), 0000023 (2022)

[4] Drukker, K., Chen, W., Gichoya, J., Gruszauskas, N., Kalpathy-Cramer, J., Koyejo, S., Myers, K., Sá, R.C., Sahiner, B., Whitney, H., et al.: Toward fairness in artificial intelligence for medical image analysis: identification and mitigation of potential biases in the roadmap from data collection to model deployment. Journal of Medical Imaging 10(6), 061104–061104 (2023)

[5] Larrazabal, A.J., Nieto, N., Peterson, V., Milone, D.H., Ferrante, E.: Gender imbalance in medical imaging datasets produces biased classifiers for computer-aided diagnosis. Proceedings of the National Academy of Sciences 117(23), 12592–12594 (2020)

[6] Seyyed-Kalantari, L., Zhang, H., McDermott, M.B., Chen, I.Y., Ghassemi, M.: Underdiagnosis bias of artificial intelligence algorithms applied to chest radiographs in under-served patient populations. Nature medicine 27(12), 2176–2182 (2021)

[7] Purushotham, S., Meng, C., Che, Z., Liu, Y.: Benchmarking deep learning models on large healthcare datasets. Journal of biomedical informatics 83, 112–134 (2018)

[8] Brain, D., Webb, G.I.: The need for low bias algorithms in classification learning from large data sets. In: European Conference on Principles of Data Mining and Knowledge Discovery, pp. 62–73 (2002). Springer

[9] Kearns, M., Neel, S., Roth, A., Wu, Z.S.: An empirical study of rich subgroup fairness for machine learning. In: Proceedings of the Conference on Fairness, Accountability, and Transparency, pp. 100–109 (2019)

[10] Huang, Y., Guo, J., Chen, W.-H., Lin, H.-Y., Tang, H., Wang, F., Xu, H., Bian, J.: A scoping review of fair machine learning techniques when using real-world data. Journal of Biomedical Informatics, 104622 (2024)

[11] Seyyed-Kalantari, L., Liu, G., McDermott, M., Chen, I.Y., Ghassemi, M.: Chexclusion: Fairness gaps in deep chest x-ray classifiers. In: BIOCOMPUTING 2021: Proceedings of the Pacific Symposium, pp. 232–243 (2020). World Scientific

[12] Sheller, M.J., Edwards, B., Reina, G.A., Martin, J., Pati, S., Kotrotsou, A., Milchenko, M., Xu, W., Marcus, D., Colen, R.R., et al.: Federated learning in medicine: facilitating multi-institutional collaborations without sharing patient data. Scientific reports 10(1), 12598 (2020)

[13] European Parliament, Council of the European Union: Regulation (EU) 2016/679 of the European Parliament and of the Council. https://data.europa.eu/eli/reg/2016/679/oj Accessed 2024-02-14

[14] European Parliament, Council of the European Union: Regulation (EU) 2016/679 of the European Parliament and of the Council. https://data.europa.eu/eli/reg/2016/679/oj Accessed 202 4-02-14

[15] Bukaty, P.: The California Consumer Privacy Act (CCPA): An Implementation Guide. IT Governance Publishing, ??? (2019). http://www.jstor.org/stable/j.ctvjghvnn Accessed 2024-02-14

[16] Pati, S., Baid, U., Edwards, B., Sheller, M., Wang, S.-H., Reina, G.A., Foley, P., Gruzdev, A., Karkada, D., Davatzikos, C., et al.: Federated learning enables big data for rare cancer boundary detection. Nature communications 13(1), 7346 (2022)

[17] Drichel, A., Holmes, B., Brandt, J., Meyer, U.: The more, the better: A study on collaborative machine learning for dga detection. In: Proceedings of the 3rd Workshop on Cyber-Security Arms Race, pp. 1–12 (2021)

[18] Pessach, D., Tassa, T., Shmueli, E.: Fairness-driven private collaborative machine learning. arXiv preprint 2109.14376 (2021)

[19] Usynin, D., Ziller, A., Makowski, M., Braren, R., Rueckert, D., Glocker, B., Kaissis, G., Passerat-Palmbach, J.: Adversarial interference and its mitigations in privacy-preserving collaborative machine learning. Nature Machine Intelligence 3(9), 749–758 (2021)

[20] McMahan, H.B., Moore, E., Ramage, D., Hampson, S., Arcas, B.: Communication-efficient learning of deep networks from decentralized data. arxiv. arXiv preprint 1602.05629 (2016)

[21] Rieke, N., Hancox, J., Li, W., Milletari, F., Roth, H.R., Albarqouni, S., Bakas, S., Galtier, M.N., Landman, B.A., Maier-Hein, K., et al.: The future of digital health with federated learning. NPJ digital medicine 3(1), 119 (2020)

[22] Patel, M., Dayan, I., Fishman, E.K., Flores, M., Gilbert, F.J., Guindy, M., Koay, E.J., Rosenthal, M., Roth, H.R., Linguraru, M.G.: Accelerating artificial intelligence: How federated learning can protect privacy, facilitate collaboration, and improve outcomes. Health informatics journal 29(4), 14604582231207744 (2023)

[23] Kairouz, P., McMahan, H.B., Avent, B., Bellet, A., Bennis, M., Bhagoji, A.N., Bonawitz, K., Charles, Z., Cormode, G., Cummings, R., et al.: Advances and open problems in federated learning. Foundations and Trends® in Machine Learning 14(1–2), 1–210 (2021)

[24] Joshi, M., Pal, A., Sankarasubbu, M.: Federated learning for healthcare domain-pipeline, applications and challenges. ACM Transactions on Computing for Healthcare 3(4), 1–36 (2022)

[25] Guha, N., Talwalkar, A., Smith, V.: One-shot federated learning. arXiv preprint 1902.11175 (2019)

[26] Wang, H., Yurochkin, M., Sun, Y., Papailiopoulos, D., Khazaeni, Y.: Federated learning with matched averaging. arXiv preprint 2002.06440 (2020)

[27] Innocenti, L., Antonelli, M., Cremonesi, F., Sarhan, K., Granados, A., Goh, V., Ourselin, S., Lorenzi, M.: Benchmarking collaborative learning methods cost-effectiveness for prostate segmentation. arXiv preprint 2309.17097 (2023)

[28] Dang, T., et al.: Weighted ensemble of deep learning models based on comprehensive learning particle swarm optimization for medical image segmentation. In: 2021 IEEE Congress on Evolutionary Computation (CEC) (2021)

[29] Dang, T., et al.: Ensemble of deep learning models with surrogate-based optimization for medical image segmentation. In: 2022 IEEE Congress on Evolutionary Computation (CEC) (2022)

[30] Kumar, A., Kim, J., Lyndon, D., Fulham, M., Feng, D.: An ensemble of finetuned convolutional neural networks for medical image classification. IEEE journal of biomedical and health informatics 21(1), 31–40 (2016)

[31] Müller, D., Soto-Rey, I., Kramer, F.: An analysis on ensemble learning optimized medical image classification with deep convolutional neural networks. Ieee Access 10, 66467–66480 (2022)

[32] Shahin, A.H., Kamal, A., Elattar, M.A.: Deep ensemble learning for skin lesion classification from dermoscopic images. In: 2018 9th Cairo International Biomedical Engineering Conference (CIBEC), pp. 150–153 (2018). IEEE

[33] Nanni, L., Ghidoni, S., Brahnam, S.: Ensemble of convolutional neural networks for bioimage classification. Applied Computing and Informatics 17(1), 19–35 (2021)

[34] Schaefer, G., Krawczyk, B., Celebi, M.E., Iyatomi, H.: An ensemble classification approach for melanoma diagnosis. Memetic Computing 6, 233–240 (2014)

[35] Xie, F., Fan, H., Li, Y., Jiang, Z., Meng, R., Bovik, A.: Melanoma classification on dermoscopy images using a neural network ensemble model. IEEE transactions on medical imaging 36(3), 849–858 (2016)

[36] De Oliveira, H., Prodel, M., Augusto, V.: Binary classification on french hospital data: benchmark of 7 machine learning algorithms. In: 2018 IEEE International Conference on Systems, Man, and Cybernetics (SMC), pp. 1743–1748 (2018). IEEE

[37] Shamshirband, S., Fathi, M., Dehzangi, A., Chronopoulos, A.T., Alinejad-Rokny, H.: A review on deep learning approaches in healthcare systems: Taxonomies, challenges, and open issues. Journal of Biomedical Informatics 113, 103627 (2021)

[38] Yèche, H., Kuznetsova, R., Zimmermann, M., Hüser, M., Lyu, X., Faltys, M., Ratsch, G.: Hirid-icu-benchmark—a comprehensive machine learning benchmark on high-resolution icu data. In: Thirty-fifth Conference on Neural Information Processing Systems Datasets and Benchmarks Track (Round 1) (2021)

[39] Olson, R.S., La Cava, W., Orzechowski, P., Urbanowicz, R.J., Moore, J.H.: Pmlb: a large benchmark suite for machine learning evaluation and comparison. BioData mining 10, 1–13 (2017)

[40] Karargyris, A., Umeton, R., Sheller, M.J., Aristizabal, A., George, J., Bala, S., Beutel, D.J., Bittorf, V., Chaudhari, A., Chowdhury, A., et al.: Medperf: open benchmarking platform for medical artificial intelligence using federated evaluation. arXiv preprint 2110.01406 (2021)

[41] Lee, G.H., Shin, S.-Y.: Federated learning on clinical benchmark data: performance assessment. Journal of medical Internet research 22(10), 20891 (2020)

[42] Chen, D., Gao, D., Kuang, W., Li, Y., Ding, B.: pfl-bench: A comprehensive benchmark for personalized federated learning. Advances in Neural Information Processing Systems 35, 9344–9360 (2022)

[43] Salmeron, J.L., Arévalo, I., Ruiz-Celma, A.: Benchmarking federated strategies in peer-to-peer federated learning for biomedical data. Heliyon 9(6) (2023)

[44] Yang, Q., Zhang, J., Hao, W., Spell, G.P., Carin, L.: Flop: Federated learning on medical datasets using partial networks. In: Proceedings of the 27th ACM SIGKDD Conference on Knowledge Discovery & Data Mining, pp. 3845–3853 (2021)

[45] Lai, F., Dai, Y., Singapuram, S., Liu, J., Zhu, X., Madhyastha, H., Chowdhury, M.: Fedscale: Benchmarking model and system performance of federated learning at scale. In: International Conference on Machine Learning, pp. 11814–11827 (2022). PMLR

[46] Asad, M., Moustafa, A., Ito, T., Aslam, M.: Evaluating the communication efficiency in federated learning algorithms. In: 2021 IEEE 24th International Conference on Computer Supported Cooperative Work in Design (CSCWD), pp. 552–557 (2021). IEEE

[47] Tedeschini, B.C., Savazzi, S., Stoklasa, R., Barbieri, L., Stathopoulos, I., Nicoli, M., Serio, L.: Decentralized federated learning for healthcare networks: A case study on tumor segmentation. IEEE access 10, 8693–8708 (2022)

[48] Terrail, J.O.d., Ayed, S.-S., Cyffers, E., Grimberg, F., He, C., Loeb, R., Mangold, P., Marchand, T., Marfoq, O., Mushtaq, E., et al.: Flamby: Datasets and benchmarks for cross-silo federated learning in realistic healthcare settings. arXiv preprint 2210.04620 (2022)

[49] Caldas, S., Duddu, S.M.K., Wu, P., Li, T., Konečny, J., McMahan, H.B., Smith, V., Talwalkar, A.: Leaf: A benchmark for federated settings. arXiv preprint 1812.01097 (2018)

[50] Chaudhari, H., Jagielski, M., Oprea, A.: Safenet: The unreasonable effectiveness of ensembles in private collaborative learning. In: 2023 IEEE Conference on Secure and Trustworthy Machine Learning (SaTML), pp. 176–196 (2023). IEEE

[51] Gupta, S., Kumar, S., Chang, K., Lu, C., Singh, P., Kalpathy-Cramer, J.: Collaborative privacy-preserving approaches for distributed deep learning using multi-institutional data. RadioGraphics 43(4), 220107 (2023)

[52] Li, T., Sahu, A.K., Zaheer, M., Sanjabi, M., Talwalkar, A., Smith, V.: Federated Optimization in Heterogeneous Networks. Proceedings of the 1 st Adaptive & Multitask Learning Workshop, Long Beach, California, 2019, 1–28 (2018) 1812.06127

[53] Karimireddy, S.P., Kale, S., Mohri, M., Reddi, S., Stich, S., Suresh, A.T.: Scaffold: Stochastic controlled averaging for federated learning. In: International Conference on Machine Learning, pp. 5132–5143 (2020). PMLR

[54] Reddi, S., Charles, Z., Zaheer, M., Garrett, Z., Rush, K., Konečny, J., Kumar, S., McMahan, H.B.: Adaptive federated optimization. arXiv preprint 2003.00295 (2020)

[55] McMahan, B., Moore, E., Ramage, D., Hampson, S., Arcas, B.A.: Communication-Efficient Learning of Deep Networks from Decentralized Data. In: ICML 2017 (2017)

[56] Safdar, K., Akbar, S., Shoukat, A.: A majority voting based ensemble approach of deep learning classifiers for automated melanoma detection. In: 2021 International Conference on Innovative Computing (ICIC), pp. 1–6 (2021). IEEE

[57] Warfield, S.K., Zou, K.H., Wells, W.M.: Simultaneous truth and performance level estimation (staple): an algorithm for the validation of image segmentation. IEEE transactions on medical imaging 23(7), 903–921 (2004)

[58] Ruta, D., Gabrys, B.: An overview of classifier fusion methods. Computing and Information systems 7(1), 1–10 (2000)

[59] Audibert, J., Michiardi, P., Guyard, F., Marti, S., Zuluaga, M.A.: Usad: Unsupervised anomaly detection on multivariate time series. In: Proceedings of the 26th ACM SIGKDD International Conference on Knowledge Discovery & Data Mining, pp. 3395–3404 (2020)

[60] Fraboni, Y., Van Waerebeke, M., Scaman, K., Vidal, R., Kameni, L., Lorenzi, M.: Sequential informed federated unlearning: Efficient and provable client unlearning in federated optimization. arXiv preprint 2211.11656 (2022)

[61] Munir, M.T., Saeed, M.M., Ali, M., Qazi, Z.A., Raza, A.A., Qazi, I.A.: Learning fast and slow: Towards inclusive federated learning. In: Joint European Conference on Machine Learning and Knowledge Discovery in Databases, pp. 384–401 (2023). Springer

[62] Nguyen, J., Malik, K., Zhan, H., Yousefpour, A., Rabbat, M., Malek, M., Huba, D.: Federated learning with buffered asynchronous aggregation. In: International Conference on Artificial Intelligence and Statistics, pp. 3581–3607 (2022). PMLR

[63] Chen, R., Yu, J.: An improved bagging neural network ensemble algorithm and its application. In: Third International Conference on Natural Computation (ICNC 2007), vol. 5, pp. 730–734 (2007). IEEE

[64] Yang, L.: Classifiers selection for ensemble learning based on accuracy and diversity. Procedia Engineering 15, 4266–4270 (2011)

[65] Yin, X., Zhu, Y., Hu, J.: A comprehensive survey of privacy-preserving federated learning: A taxonomy, review, and future directions. ACM Computing Surveys (CSUR) 54(6), 1–36 (2021)

[66] Bak, M., Madai, V.I., Celi, L.A., Kaissis, G.A., Cornet, R., Maris, M., Rueckert, D., Buyx, A., McLennan, S.: Federated learning is not a cure-all for data ethics. Nature Machine Intelligence, 1–3 (2024)

[67] Hu, L., Evans, D.: Secure aggregation for wireless networks. In: 2003 Symposium on Applications and the Internet Workshops, 2003. Proceedings., pp. 384–391 (2003). IEEE

[68] Mansouri, M., Onen, M., Jaballah, W.B., Conti, M.: Sok: Secure aggregation based on cryptographic schemes for federated learning. Proc. Priv. Enhancing Technol, 140–157 (2023)

[69] Wei, K., Li, J., Ding, M., Ma, C., Yang, H.H., Farokhi, F., Jin, S., Quek, T.Q., Poor, H.V.: Federated learning with differential privacy: Algorithms and performance analysis. IEEE Transactions on Information Forensics and Security 15, 3454–3469 (2020)

[70] Ronneberger, O., Fischer, P., Brox, T.: U-net: Convolutional networks for biomedical image segmentation. In: MICCAI 2015 (2015). Springer

[71] Myronenko, A.: 3d mri brain tumor segmentation using autoencoder regularization. brainlesion glioma mult. scler. Stroke Trauma. Brain Inj.-BrainLes 2019, 11384 (2018)

[72] Antonelli, M., Reinke, A., Bakas, S., Farahani, K., Kopp-Schneider, A., Landman, B.A., Litjens, G., Menze, B., Ronneberger, O., Summers, R.M., et al.: The medical segmentation decathlon. Nature communications 13(1), 4128 (2022)

[73] Litjens, G., Toth, R., Van De Ven, W., Hoeks, C., Kerkstra, S., Van Ginneken, B., Vincent, G., Guillard, G., Birbeck, N., Zhang, J., et al.: Evaluation of prostate segmentation algorithms for mri: the promise12 challenge. Medical image analysis 18(2), 359–373 (2014)

[74] Armato III, S.G., Huisman, H., Drukker, K., Hadjiiski, L., Kirby, J.S., Petrick, N., Redmond, G., Giger, M.L., Cha, K., Mamonov, A., et al.: Prostatex challenges for computerized classification of prostate lesions from multiparametric magnetic resonance images. Journal of Medical Imaging 5(4), 044501–044501 (2018)

[75] Janosi, A., Steinbrunn, W., Pfisterer, M., Detrano, R.: Heart disease data set. The UCI KDD Archive (1988)

[76] team, B.: Ixi dataset. https://brain-development.org/ixi-dataset/

[77] Gutman, D., Codella, N.C., Celebi, E., Helba, B., Marchetti, M., Mishra, N., Halpern, A.: Skin lesion analysis toward melanoma detection: A challenge at the international symposium on biomedical imaging (isbi) 2016, hosted by the international skin imaging collaboration (isic). arXiv preprint 1605.01397 (2016)

[78] Tan, M., Le, Q.: Efficientnet: Rethinking model scaling for convolutional neural networks. In: International Conference on Machine Learning, pp. 6105–6114 (2019). PMLR

[79] Tomczak, K., Czerwińska, P., Wiznerowicz, M.: Review the cancer genome atlas (tcga): an immeasurable source of knowledge. Contemporary Oncology/Wsp ó lczesna Onkologia 2015(1), 68–77 (2015)

[80] Heller, N., Sathianathen, N., Kalapara, A., Walczak, E., Moore, K., Kaluzniak, H., Rosenberg, J., Blake, P., Rengel, Z., Oestreich, M., et al.: The kits19 challenge data: 300 kidney tumor cases with clinical context, ct semantic segmentations, and surgical outcomes. arXiv preprint 1904.00445 (2019)

[81] Isensee, F., Jaeger, P.F., Kohl, S.A., Petersen, J., Maier-Hein, K.H.: nnu-net: a self-configuring method for deep learning-based biomedical image segmentation. Nature methods 18(2), 203–211 (2021)

[82] Pati, S., Baid, U., Zenk, M., Edwards, B., Sheller, M., Reina, G.A., Foley, P., Gruzdev, A., Martin, J., Albarqouni, S., et al.: The federated tumor segmentation (fets) challenge. arXiv preprint 2105.05874 (2021)

[83] Kingma, D.P., Ba, J.: Adam: A method for stochastic optimization. arXiv preprint 1412.6980 (2014)

[84] Zaheer, M., Reddi, S., Sachan, D., Kale, S., Kumar, S.: Adaptive methods for 31 nonconvex optimization. Advances in neural information processing systems 31 (2018)

[85] Lydia, A., Francis, S.: Adagrad—an optimizer for stochastic gradient descent. Int. J. Inf. Comput. Sci 6(5), 566–568 (2019)

[86] Wang, L., Xu, S., Wang, X., Zhu, Q.: Addressing class imbalance in federated learning. In: Proceedings of the AAAI Conference on Artificial Intelligence, vol. 35, pp. 10165–10173 (2021)

[87] Sarkar, D., Narang, A., Rai, S.: Fed-focal loss for imbalanced data classification in federated learning. arXiv preprint 2011.06283 (2020)

[88] Pillutla, K., Kakade, S.M., Harchaoui, Z.: Robust aggregation for federated learning. IEEE Transactions on Signal Processing 70, 1142–1154 (2022)

[89] Qi, P., Chiaro, D., Guzzo, A., Ianni, M., Fortino, G., Piccialli, F.: Model aggregation techniques in federated learning: A comprehensive survey. Future Generation Computer Systems (2023)

[90] Liu, J., Wang, J.H., Rong, C., Xu, Y., Yu, T., Wang, J.: Fedpa: An adaptively partial model aggregation strategy in federated learning. Computer Networks 199, 108468 (2021)

[91] He, C., Annavaram, M., Avestimehr, S.: Group knowledge transfer: Federated learning of large cnns at the edge. Advances in Neural Information Processing Systems 33, 14068–14080 (2020)

[92] Lian, Z., Yang, Q., Wang, W., Zeng, Q., Alazab, M., Zhao, H., Su, C.: Deep-fel: Decentralized, efficient and privacy-enhanced federated edge learning for healthcare cyber physical systems. IEEE Transactions on Network Science and Engineering 9(5), 3558–3569 (2022)

[93] Hansen, L.K., Salamon, P.: Neural network ensembles. IEEE transactions on pattern analysis and machine intelligence 12(10), 993–1001 (1990)

[94] Kuncheva, L.I.: A theoretical study on six classifier fusion strategies. IEEE Transactions on pattern analysis and machine intelligence 24(2), 281–286 (2002)

[95] Ganaie, M.A., Hu, M., Malik, A., Tanveer, M., Suganthan, P.: Ensemble deep learning: A review. Engineering Applications of Artificial Intelligence 115, 105151 (2022)

[96] Adiga, A., Wang, L., Hurt, B., Peddireddy, A., Porebski, P., Venkatramanan, S., Lewis, B.L., Marathe, M.: All models are useful: Bayesian ensembling for robust high resolution covid-19 forecasting. In: Proceedings of the 27th ACM SIGKDD Conference on Knowledge Discovery & Data Mining, pp. 2505–2513 (2021)

[97] Cremonesi, F.e.a.: Fed-biomed: Open, transparent and trusted federated learning for real-world healthcare applications. arXiv preprint 2304.12012 (2023)

[98] Paszke, A., Gross, S., Massa, F., Lerer, A., Bradbury, J., Chanan, G., Killeen, T., Lin, Z., Gimelshein, N., Antiga, L., Desmaison, A., Kopf, A., Yang, E., DeVito, Z., Raison, M., Tejani, A., Chilamkurthy, S., Steiner, B., Fang, L., Bai, J., Chintala, S.: Pytorch: An imperative style, high-performance deep learning library. In: Advances in Neural Information Processing Systems 32, pp. 8024–8035. Curran Associates, Inc., ??? (2019). http://papers.neurips.cc/paper/9015-pytorch-an-imperative-style-high-performance-deep-learning-library.pdf

[99] Cardoso, M.J., Li, e.a.: MONAI: An open-source framework for deep learning in healthcare (2022) 10.48550/arXiv.2211.02701

[100] Pérez-García, F., Sparks, R., Ourselin, S.: Torchio: a python library for efficient loading, preprocessing, augmentation and patch-based sampling of medical images in deep learning. Computer Methods and Programs in Biomedicine 208, 106236 (2021)

[101] Beare, R., Lowekamp, B., Yaniv, Z.: Image segmentation, registration and characterization in r with simpleitk. Journal of statistical software 86 (2018)

[102] Yaniv, Z., Lowekamp, B.C., Johnson, H.J., Beare, R.: Simpleitk image-analysis notebooks: a collaborative environment for education and reproducible research. Journal of digital imaging 31(3), 290–303 (2018)

[103] Lowekamp, B.C., Chen, D.T., Ibáñez, L., Blezek, D.: The design of simpleitk. Frontiers in neuroinformatics 7, 45 (2013)

[104] Cuocolo, R., Stanzione, A., Castaldo, A., De Lucia, D.R., Imbriaco, M.: Quality control and whole-gland, zonal and lesion annotations for the prostatex challenge public dataset. European Journal of Radiology 138, 109647 (2021)

[105] Loshchilov, I., Hutter, F.: Decoupled weight decay regularization. In: arXiv Preprint 1711.05101 (2017)

[106] Arora, A.: Siim-isic melanoma classification - my journey to a top 5 https://amaarora.github.io/2020/08/23/siimisic.html

